# Highly contiguous reference genome assembly of the endangered Orcés’ blue whiptail *Holcosus orcesi*

**DOI:** 10.64898/2026.05.14.725226

**Authors:** Gabriela Pozo, Diego F. Cisneros-Heredia, Doménica Barragán-Orbe, Juan C. Sánchez-Nivicela, Ernesto Arbeláez, Maria de Lourdes Torres

## Abstract

*Holcosus orcesi*, the Orcés’ Blue Whiptail, is a Critically Endangered lizard endemic to the upper Jubones River basin in southern Ecuador. Restricted to a narrow elevational range within semi-arid Andean shrublands, it represents one of the few montane members of a predominantly lowland lineage. Here we present the first high-quality reference genome for *H. orcesi*, generated using Oxford Nanopore Technologies long-read sequencing. The assembly spans 1.68 Gb across only 91 contigs, with an N50 of 76.2 Mb and a BUSCO completeness of 96.8%, making it among the most contiguous and complete squamate genomes to date. Structural annotation predicted 25,682 genes, of which 85% showed homology to known proteins and 45% were assigned Gene Ontology terms. Repetitive elements accounted for 46.3% of the genome, with LINEs representing the predominant class. This genome provides a foundational resource for future evolutionary, comparative and conservation-genomic research of *H. orcesi* and other mountain reptiles, enabling studies of population genomics, local adaptation, and genomic erosion in isolated populations. By expanding the genomic representation of tropical montane reptiles, this work helps address longstanding phylogenetic and geographic gaps in global biodiversity genomics and provides a foundation for evidence-based conservation of *H. orcesi* and related taxa.

## 1. Introduction

*Holcosus orcesi* [1], commonly known as Orcés’ Blue Whiptail, is a small teiid lizard endemic to the upper Jubones River basin on the southwestern Andes of Ecuador, where it occupies a narrow elevational range between 1,000 and 1,600 m (Fig 1A and 1B) [1,2]. The species went unrecorded for nearly six decades and was presumed extinct until its rediscovery in the mid-2010s [3,4]. It inhabits xeric montane shrublands and persists in small, isolated patches of native vegetation surrounded by pastures and croplands [1,3]. The region is semi-arid, with low annual rainfall (≈400 mm/year), concentrated between January and May, followed by a prolonged dry season, and experiences mild days and cool nights [5–7]. *Holcosus orcesi* is strictly diurnal and heliothermic, active only under full sunlight and retreating when clouds obscure the sun. It forages swiftly among roots, thorny shrubs, and sun-exposed rocky slopes, feeding mainly on beetles and grasshoppers, and relies on vigilance and sprint speed as primary defenses [1].

**Figure 1.**
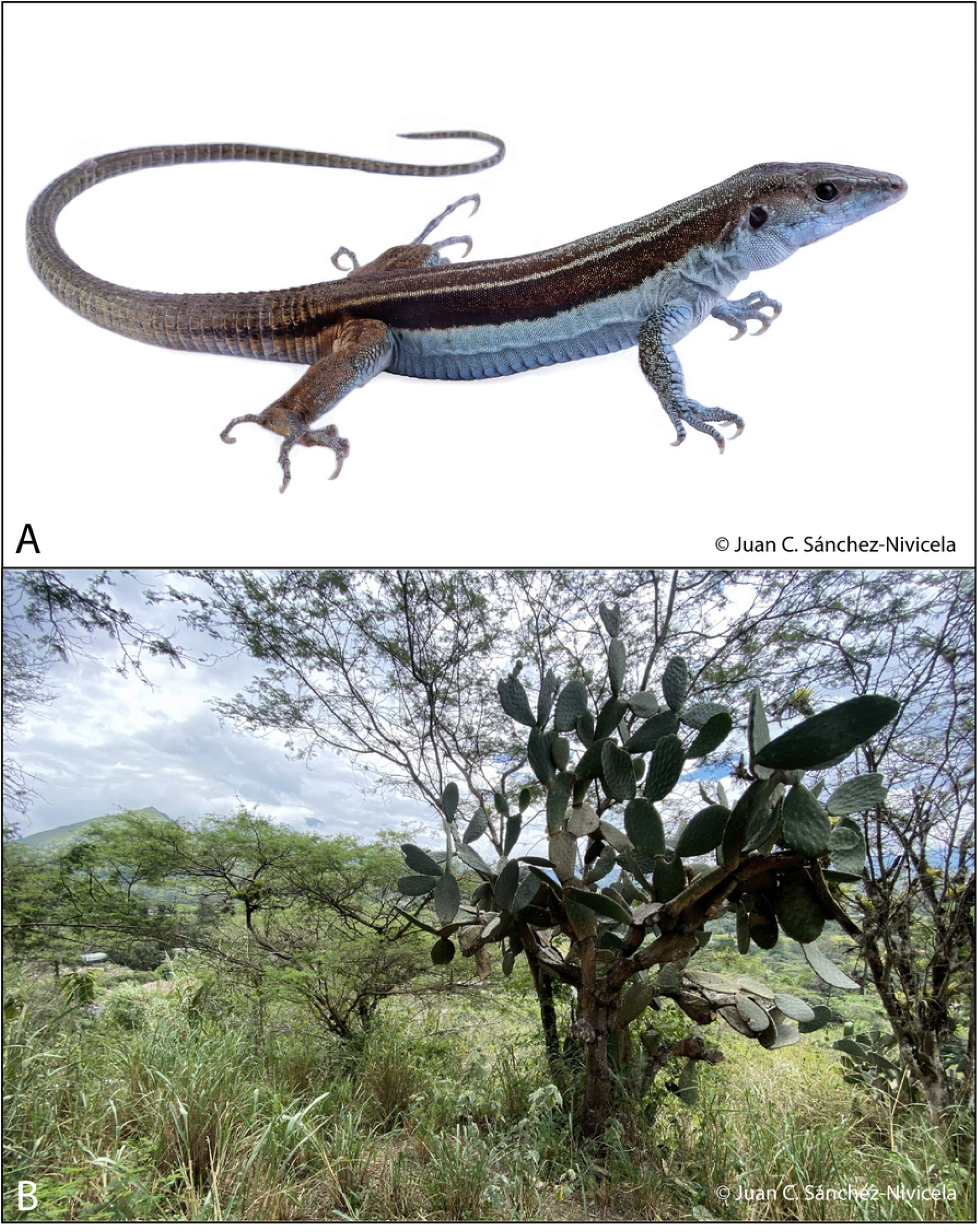
*Holcosus orcesi* and its natural habitat. Representative individual of the Ecuadorian endemic lizard *Holcosus orcesi* and its natural dry forest habitat in southern Ecuador. A) Adult *H. orcesi* individual. B) Typical habitat characterized by xeric vegetation and cacti

The genus *Holcosus* extends from southern Mexico to north-western South America, and *H. orcesi* is one of the few montane species within this predominantly lowland radiation [2]. Its apparent narrow thermal niche and dependence on open, sunlit microhabitats likely increase its sensitivity to climatic and habitat changes. Habitat conversion for agriculture, invasive predators and poaching have contributed to population decline, leading the IUCN to list it as Critically Endangered [4]. As a geographically restricted species occupying a biogeographically unusual position within its genus, *H. orcesi* exemplifies evolutionary persistence under severe environmental stress [1,2]. Studying its genome could offer a valuable opportunity to understand how this lineage has adapted to an environmentally demanding, semi-arid montane habitat. However, genomic resources for Andean species remain scarce, limiting comparative work on the evolutionary and ecological diversification of this fauna.

High-quality genomic resources have become indispensable tools for modern biodiversity conservation. Reference genomes provide the foundation for investigating species’ evolutionary histories, demographic trajectories, and adaptive potential, particularly in lineages with restricted distributions and elevated extinction risk [8,9]. In threatened taxa such as *H. orcesi*, these resources are especially valuable because they support future analyses of population structure, inbreeding, and evolutionary history that cannot be addressed fully with traditional genetic markers alone. The availability of a reference genome opens the possibility of identifying genetic signatures of adaptation to its harsh high-altitude environment, including tolerance to hypoxia, UV exposure, and cold temperatures. Genomic approaches have revealed high-altitude adaptations in other vertebrates, including hypoxia-related pathways in yak (*Bos grunniens*) [10], the Tibetan ground tit (*Parus humilis*) [11], and multiple mammals such as the Tibetan sheep and plateau zokor [12,13].

The integration of genomics into conservation has already transformed our ability to monitor population health. Genomic data enable more accurate estimates of effective population size, levels of genetic diversity, and the genomic basis of inbreeding depression, which are crucial to designing effective management plans [9,14]. In particular, in endangered species with small and isolated populations, such as *H. orcesi*, these resources allow detection of deleterious alleles and runs of homozygosity that may otherwise go unnoticed through traditional genetic markers [14–16].

Beyond species-specific applications, genomic resources contribute to comparative and macroevolutionary studies. Initiatives such as the Earth BioGenome Project and related efforts have emphasized the importance of expanding genomic representation across taxonomic and geographic scales, noting significant biases in available references toward temperate species [8,17]. By contributing the genome of a narrowly distributed Andean reptile, our study helps to fill this latitudinal and phylogenetic gap, providing a resource for Neotropical squamates that benefits both regional and global biodiversity knowledge.

Ultimately, genomic resources represent one of the most powerful tools available to address the global loss of biodiversity. They inform not only the immediate conservation of threatened taxa, but also broader strategies aimed at preserving adaptive capacity in the face of rapid environmental change [18]. In this context, the genome of *H. orcesi* will serve as a cornerstone for future studies in evolutionary biology, population genomics, and conservation management of Andean reptiles. Here, we present the first reference genome assembly for *Holcosus orcesi*, generated using Oxford Nanopore long-read sequencing, together with structural annotation and repeat characterization. This resource will serve as a cornerstone for future studies in evolutionary biology, population genomics, and conservation management of Andean reptiles, and supports research on *H. orcesi, Holcosus*, and other poorly represented tropical montane squamates.

## 2. Materials and Methods

### 2.1 Sample Collection and DNA Extraction

A single specimen of *Holcosus orcesi* was collected in February 2025 from the upper Jubones River basin, southwestern Ecuador. The exact locality is withheld to reduce poaching but is available upon reasonable request from the Museo de Zoología, Universidad San Francisco de Quito. The individual was captured by hand, and a licensed wildlife veterinarian collected a blood sample. The voucher specimen is deposited at the Museo de Zoología, Universidad San Francisco de Quito, Ecuador (ZSFQ). Whole blood was preserved in K2-EDTA and stored at –80 °C until DNA extraction. All procedures followed the guidelines for use of live reptiles in field research [19]. Research activities were conducted under authorization from the Ministerio del Ambiente, Agua y Transición Ecológica del Ecuador (research permit No. MAATE-ARSFC-2025-0144 and framework contract for access to genetic resources No. MAATE-DBI-CM-2023-0313).

Ultra-high molecular weight genomic DNA was isolated from blood obtained from this same individual using two protocols: the Monarch® HMW DNA Extraction Kit (T3050) for nucleated blood, and the Monarch® Genomic DNA Purification Kit (T3010) for nucleated red blood cells. For HMW extraction, 15 μl of blood was used as input material, while 10 μl was used for genomic DNA extraction. In both cases DNA was eluted in 100 μl. DNA integrity was assessed by electrophoresis on a 1.5% agarose gel, followed by quantification using a Qubit 4 Fluorometer.

### 2.2 Library Preparation and Sequencing

Two sequencing libraries were prepared using the Ligation Sequencing Kit SQK-LSK114 (Oxford Nanopore Technologies), starting with 1000 ng of input DNA. The library preparation protocol was followed with minor modifications in the DNA repair and end-prep step (extended incubation time of 15 minutes at 20°C and 15 minutes at 65°C and a single ethanol wash), and in the adapter ligation and clean-up step (extended incubation time of 20 minutes at room temperature and a single Long Fragment Buffer wash).

Sequencing was performed on a PromethION 2 Solo device using two R10.4.1 flow cells, each run for ∼21 hours. Data acquisition and run control were managed with MinKNOW (v6.5.14), and raw signals were base called with Dorado (v7.9.8) in high-accuracy mode, using a minimum Q score threshold of 7.

### 2.3 Read Processing, Genome Assembly and Quality Assessment

All passed Oxford Nanopore reads with quality scores ≥7 were concatenated, and sequencing adapters were trimmed using Porechop v0.2.3_seqan2.1.1 [20]. Read quality and length distributions were first assessed with NanoPlot v1.32.1 [21]. Reads shorter than 5,000 bp were discarded with NanoFilt v2.3.0 prior to assembly [21]. The filtered dataset was assembled de novo using Hifiasm v0.25.0-r726 [22] in ONT-optimized mode, and primary contigs were extracted from the assembly graph with gfatools v0.4-r214-dirty [23]. To improve consensus accuracy, raw ONT reads were mapped back to the draft assembly with minimap2 v2.30-r1287 [24], and the resulting alignments were used to polish the genome with Racon v1.4.20 [25]. Two successive rounds of Racon polishing were performed to maximize assembly accuracy.

Assembly completeness was quantified with BUSCO v5.8.3 [26] (mode: genome) using the squamata_odb12 lineage dataset, which surveys 11,294 near-universal single-copy orthologs expected to be present in any squamate genome (Manni et al., 2021). To assess contiguity, we ran QUAST v5.0.2 [27] with default settings to report standard metrics, including total assembly length, number of contigs (≥500 bp), largest contig, N50/L50, N90/L90, and GC content.

### 2.4 Genome Annotation and Identification of Repetitive Elements

Structural annotation was performed in OmicsBox v3.4.5 (BioBam Bioinformatics, Valencia, Spain) using AUGUSTUS, with *Gallus gallus* specified as the closest available model species. To increase lineage-specific accuracy, protein evidence from members of the Teiidae family was provided as external support for gene model construction. Prior to annotation, the genome was soft masked with the Dfam profile HMM library via HMMER.

Functional annotation of the predicted gene set was carried out in OmicsBox through two complementary approaches. First, InterProScan v5.72-103.0 was used to identify conserved protein domains, families, and motifs, and to assign Gene Ontology (GO) terms [28]. Second, similarity-based annotation was performed with DIAMOND BLAST against the NCBI non-redundant (nr) protein database, enabling functional assignments based on homology to previously characterized proteins [29]. The combined evidence from domain analysis and sequence similarity provided robust functional characterization of the annotated gene models.

Repetitive elements were identified and categorized using RepeatModeler2 (v2.0.7) [30]. First, a custom repeat library was constructed from the polished genome assembly. This library was then used by RepeatMasker (v4.1.9) to annotate and soft-mask repetitive elements in the genome [31].

## 3. Results

### 3.1 Sequencing Output and Assembly Statistics

A total of 10.11 million reads were generated using Oxford Nanopore sequencing, yielding 79.03 Gb of long-read data. All passed reads had a quality score greater than 7, with a mean read quality of 14.9. Regarding read size, the average read length was 7.8 Kb, with an N50 read length of 17.8 Kb. The coverage (∼49x) was calculated based on the expected genome size from a closely related species, *Holcosus undulatus*, which is 1.6 Gb [32] (Table 1).

**Table 1.** Sequencing output summary.

| Generated bases | Mean read length | Mean read quality | Read length N50 | Coverage |
| --- | --- | --- | --- | --- |
| 79.03 Gb | 7.8 kb | 14.9 | 17.8 kb | 49x |

The final assembly of *H. orcesi* generated with Hifiasm spans 1.68 Gb, with an N50 of 76.2 Mb. The assembly is divided into 91 contigs, with 8 contigs representing 50% of the genome (L50), and the largest contig measuring 179.6 Mb. In terms of completeness, BUSCO analysis recovered 96.8 % of complete orthologs (10,939 BUSCOs), including 96.5% single-copy and 0.4 % duplicated genes. Additionally, 1.4% of the genes were classified as fragmented and 1.7% as missing (Table 2, Fig 2).

**Table 2.** Assembly metrics for the *Holcosus orcesi* polished genome.

| Parameter | Value |
| --- | --- |
| Total length (bp) | 1,676,717,785 |
| # contigs | 91 |
| Largest contig (bp) | 179,640,633 |
| N50 (bp) | 76,215,922 |
| L50 | 8 |
| GC (%) | 42.39 % |
| Complete BUSCOs | 96.8 % (10,936) |
| Single copy | 96.5 % (10,897) |
| Duplicated | 0.4 % (42) |
| Fragmented BUSCOs | 1.4 % (159) |
| Missing BUSCOs | 1.7 % (196) |

**Figure 2.**
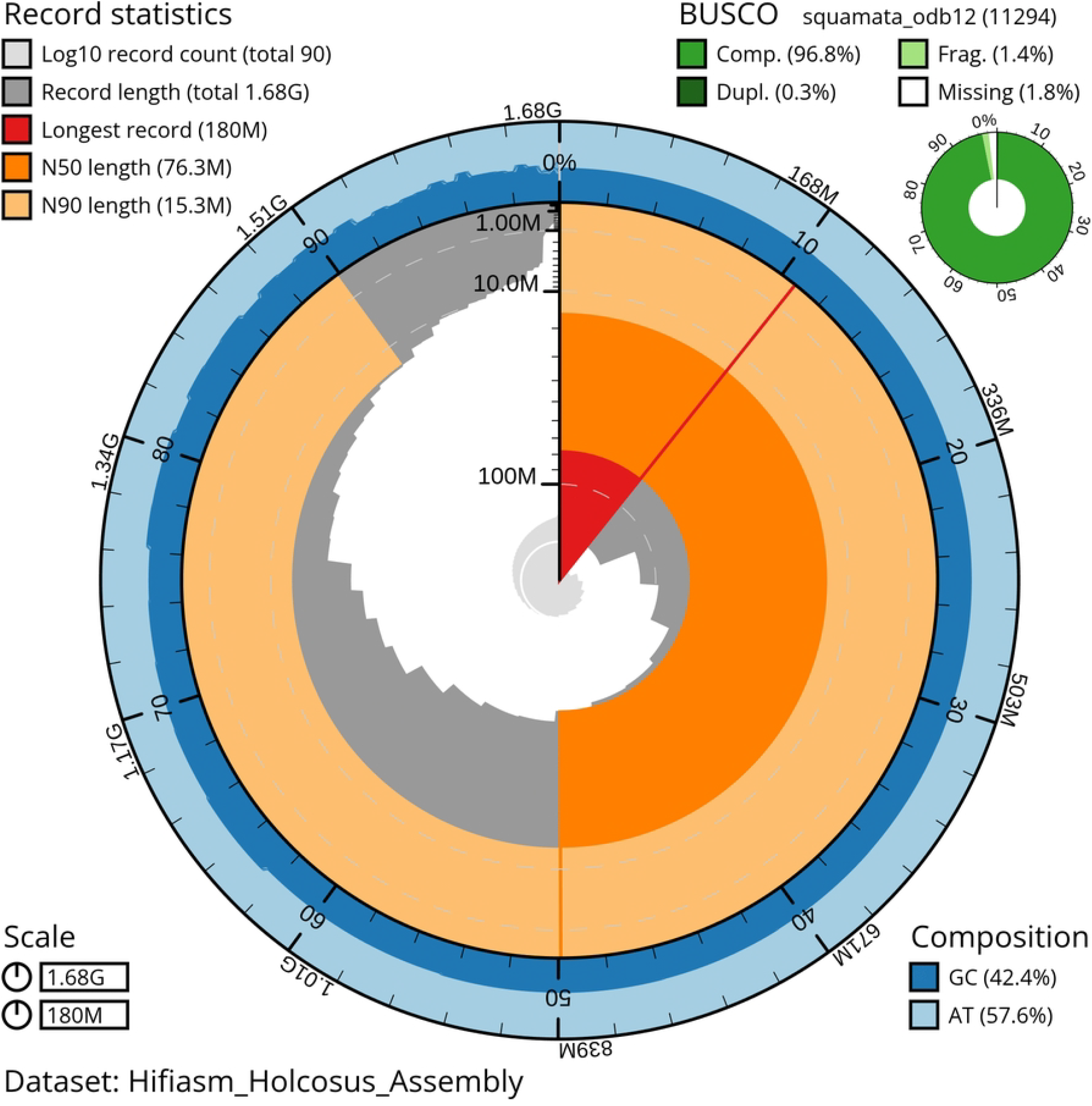
Reference genome assembly statistics for *Holcosus orcesi*. Snailplot summary statistics of the *Holcosus orcesi* reference genome assembly generated using Oxford Nanopore sequencing. The plot summarizes assembly size, contiguity metrics, GC/AT composition, and BUSCO completeness based on the squamata_odb12 dataset

### 3.2 Genome Annotation and Repetitive Elements

The final gene set consisted of 25,682 predicted genes, with a total of 152,998 exons and 127,316 introns. On average, each gene encoded a single mRNA, containing 5.9 exons and 4.9 introns. The mean gene length was 18,052 bp, with exons averaging 244 bp and introns 3,347 bp in length (Table 3). In this gene set, 84.92% of proteins showed significant similarity to known proteins in the nr database, 85.09% contained at least one conserved domain, and 44.79% were assigned at least one Gene Ontology (GO) term (Table 4).

**Table 3.** Genome annotation statistics.

| Statistic | Value |
| --- | --- |
| Number of gene | 25,682 |
| Number of exons | 152,998 |
| Number of introns in CDS | 127,316 |
| Overlapping genes | 0 |
| Mean mRNAs per gene | 1 |
| Mean exons per mRNA | 5.9 |
| Mean introns per mRNA | 4.9 |
| Mean gene length (bp) | 18,052 |
| Mean exon length (bp) | 244 |
| Mean intron length (bp) | 3,347 |

**Table 4.** Functional annotation of the predicted gene set.

| Annotation step | Metric | Value | Percentage of predicted gene set |
| --- | --- | --- | --- |
| InterProScan | Proteins with $\geq 1$ conserved domain | 21,853 | 85.09% |
| DIAMOND BLAST (nr) | Proteins with significant hit (e-value $< 1e-5$) | 21,811 | 84.92% |
| Gene Ontology | GO mapped | 13,952 | 54.33% |
|  | GO annotated | 11,502 | 44.79% |

InterProScan identified conserved domains and families in 21,853 proteins, representing 85.1% of the predicted gene set. Among PFAM domains, the most frequent were reverse transcriptase (5.1%), integrase catalytic core (3.7%), GPCR family 3 C-terminal (2.4%), zinc finger C2H2-type (1.9%), and protein kinase domains (1.5%). Similarly, the most represented InterPro families were the ribonuclease H superfamily (7.7%), DNA/RNA polymerase superfamily (7.5%), ribonuclease H-like superfamily (5.3%), reverse transcriptase/diguanylate cyclase domain family (4.4%), and the L1 transposable element C-terminal domain (4.4%). These patterns reflect the prevalence of mobile element-related proteins, signaling components, and nucleic acid–binding factors in the *H. orcesi* proteome (Table 5).

**Table 5.**
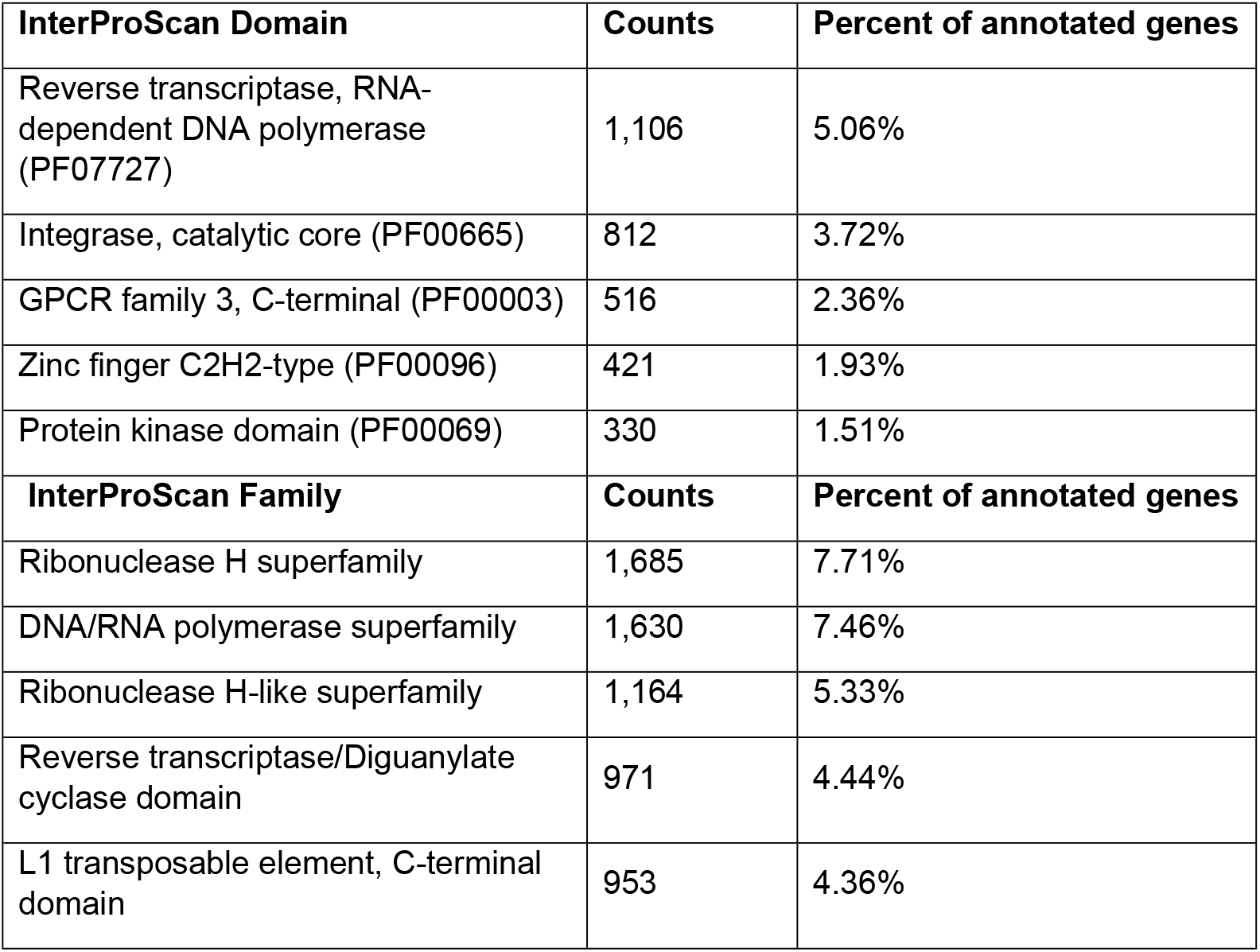
Top InterProScan Domains and Families.

| InterProScan Domain | Counts | Percent of annotated genes |
| --- | --- | --- |
| Reverse transcriptase, RNA-dependent DNA polymerase (PF07727) | 1,106 | 5.06% |
| Integrase, catalytic core (PF00665) | 812 | 3.72% |
| GPCR family 3, C-terminal (PF00003) | 516 | 2.36% |
| Zinc finger C2H2-type (PF00096) | 421 | 1.93% |
| Protein kinase domain (PF00069) | 330 | 1.51% |
| InterProScan Family | Counts | Percent of annotated genes |
| Ribonuclease H superfamily | 1,685 | 7.71% |
| DNA/RNA polymerase superfamily | 1,630 | 7.46% |
| Ribonuclease H-like superfamily | 1,164 | 5.33% |
| Reverse transcriptase/Diguanylate cyclase domain | 971 | 4.44% |
| L1 transposable element, C-terminal domain | 953 | 4.36% |

Gene Ontology (GO) analysis assigned functional categories to 11,544 proteins (44.8% of the predicted gene set). Within the Biological Process domain, the most represented categories were cellular processes (7.5%), metabolic processes (3.9%), and regulation of biological processes (3.9%), highlighting the prevalence of genes involved in fundamental cellular and regulatory pathways. In the Molecular Function domain, the majority of proteins were associated with binding activities. This included protein, nucleic acid, and small molecule binding (together accounting for more than 10% of annotated proteins), followed by catalytic activity, particularly hydrolase and transferase functions.

For the Cellular Component domain, proteins were primarily assigned to intracellular anatomical structures (e.g., organelles, cytoplasm) and to membrane-associated compartments, reflecting the structural and functional organization of the *H. orcesi* proteome (Fig 3).

**Figure 3.**
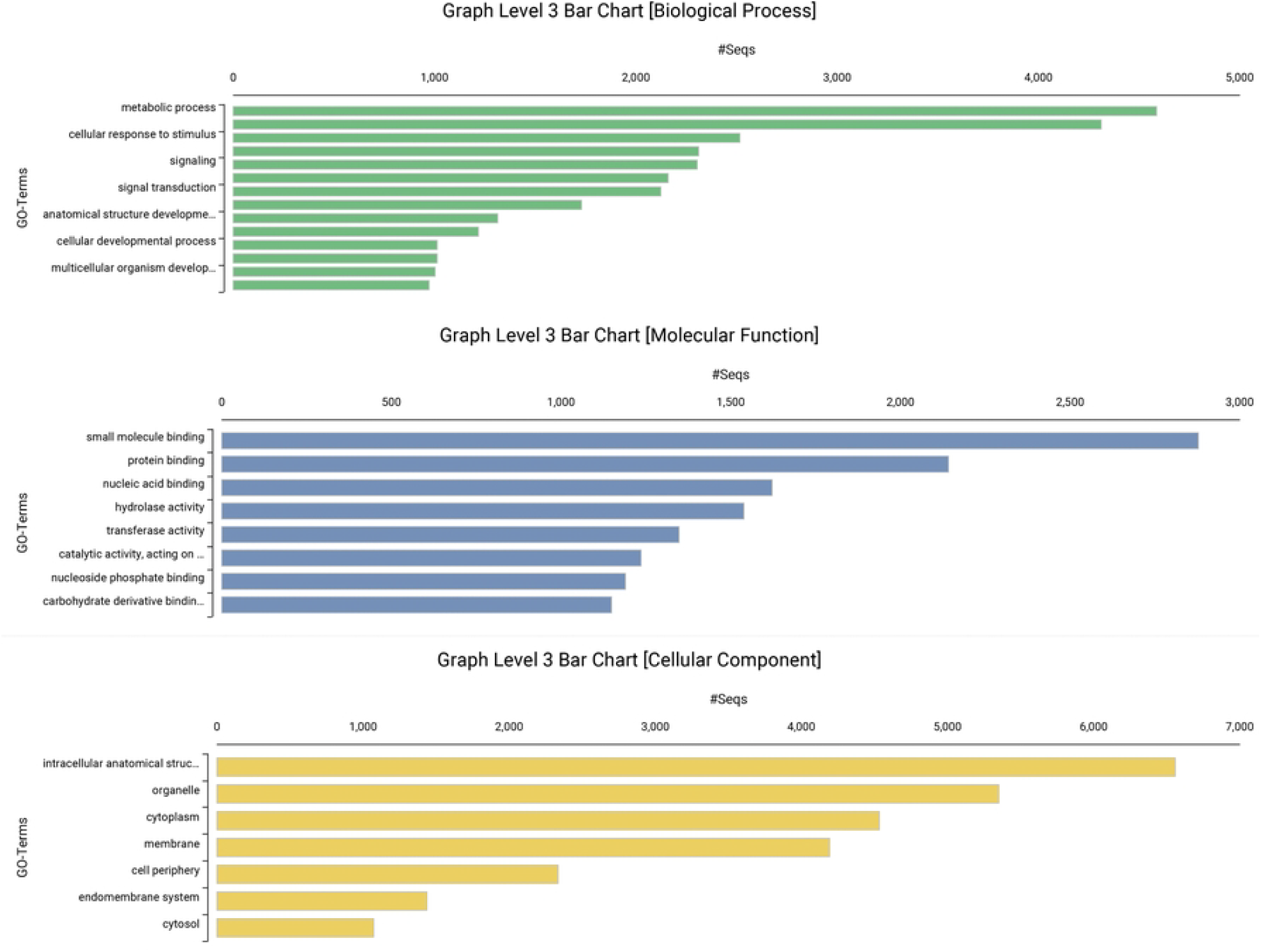
Gene Ontology Level 3 functional annotation of the Holcosus orcesi genome. Distribution of Gene Ontology (GO) Level 3 functional annotations for the annotated genes of *Holcosus orcesi*. Functional categories are grouped into the three principal GO domains: Biological Process, Molecular Function, and Cellular Component.

Analysis of the *Holcosus orcesi* genome assembly revealed that repetitive elements occupy approximately 46.29% of the total genome sequence. Among these, retroelements were the most abundant class, comprising 19.05% of the genome and including SINEs (1.45%), LINEs (13.73%), and LTR elements (3.87%). LINEs were the most prevalent subclass within retroelements, both in terms of number (960,488 elements) and sequence coverage (230,443,544 bp) (Table 6).

**Table 6.** Repetitive elements report for the *H. orcesi* genome.

| Repetitive elements | Number of elements | Length occupied (bp) | % of sequence |
| --- | --- | --- | --- |
| Retroelements | 1,411,885 | 319,681,646 | 19.05 % |
| SINEs | 181,916 | 24,258,473 | 1.45 % |
| LINEs | 960,488 | 230,443,544 | 13.73 % |
| LTR elements | 269,481 | 64,979,629 | 3.87 % |
| DNA transposons | 11,995 | 3,780,565 | 0.23 % |
| Unclassified | 2,470,382 | 419,569,037 | 25.01 % |
| Small RNA | 49,987 | 5,194,979 | 0.31 % |
| Simple repeats | 345,672 | 24,472,508 | 1.46 % |
| Low complexity | 36,658 | 4,007,354 | 0.24 % |

DNA transposons were relatively scarce, representing only 0.23% of the genome. In contrast, unclassified repetitive elements occupied the largest proportion of the genome at 25.01%, indicating a significant portion of repeats that could not be assigned to known categories. Other repeat types included small RNAs (0.31%), simple repeats (1.46%), and low complexity regions (0.24%) (Table 6).

## 4. Discussion

The *Holcosus orcesi* genome assembly presented in this paper is of very high quality, both in terms of completeness and contiguity. The assembly spans 1.68 Gb in only 91 contigs, with an N50 of 76.2 Mb and a maximum contig length of 179 Mb. BUSCO analysis using the squamata_odb12 dataset (11,294 near-universal single-copy orthologs) recovered 96.8% complete genes, indicating an excellent representation of the expected gene number [33,34]. This combination of high contiguity and completeness provides a robust foundation for downstream analyses, including the identification of structural variants, population genomic studies, and functional analyses of adaptive traits. In particular, the recovery of nearly all conserved orthologs ensures that comparative genomics and annotation pipelines can be performed without the biases introduced by missing gene families. Such metrics indicate that the *H. orcesi* genome meets current standards for reference-quality assemblies in non-model reptiles [26,35].

Within the genus, the only other available reference is for *H. undulatus* (GCA_046270765.1), which spans ∼1.6 Gb but is highly fragmented across more than 800,000 scaffolds. This contrast underscores the advances achieved in our study, where contig number was reduced by more than four orders of magnitude while achieving greater contiguity and completeness, placing it within the upper range of current lizard assemblies [34,36]. Compared to other squamate assemblies, our results are on par with recent chromosome-level references for well-studied lizards such as *Anolis carolinensis* and *Sceloporus undulatus*, where N50 values often exceed 10–50 Mb [33,37]. Thus, the *H. orcesi* genome not only represents a substantial improvement within its genus but also provides a genomic resource of comparable quality to leading reptile references, filling a critical gap in representation of high-Andean reptiles.

The structural annotation values (Table 3) are consistent with those reported for other squamate genomes. For example, the green anole (*Anolis carolinensis*) genome contains ∼19,000 protein-coding genes [37], while the Argentine black and white tegu (*Salvator merianae*) and other lizard genomes typically report between 20,000 and 26,000 predicted genes [38]. The average exon and intron lengths observed here also fall within the range described for reptiles, where exons are generally short (<300 bp) and introns considerably longer (>2 kb) [39]. Thus, the structural annotation of *Holcosus orcesi* is broadly comparable to other well-assembled squamate genomes, supporting the reliability of the predicted gene models.

The functional annotation revealed that the most enriched domain families in the *H. orcesi* proteome were strongly associated with retroelements, including reverse transcriptase, integrase, and ribonuclease H families. This reflects the pervasive influence of mobile elements on the squamate gene space, as observed in other reptilian annotations [40,41]. Nevertheless, functional categories were broadly distributed across core cellular and metabolic pathways. GO analysis revealed a predominance of genes involved in cellular and metabolic processes, binding functions (protein, nucleic acid, and small molecules), and catalytic activity, mirroring the general functional composition observed in other reptiles [42]. The assignment of ∼45% of predicted proteins to GO terms aligns with annotation levels reported in other non-model squamates, where limited experimental evidence and incomplete functional databases constrain full annotation [37,41].

The repetitive landscape of the *H. orcesi* genome is consistent with the high repeat content typically found in squamates, with ∼47% of the genome annotated as repetitive, including substantial contributions from LINEs (∼14%) and LTR retrotransposons (∼4%) [40,43,44]. This proportion falls within the broad range reported across lizards and snakes, where repeat content varies strikingly from ∼25% to over 70% despite relatively constrained genome sizes [40,43,44]. Such variation contrasts with the more conserved repeat landscapes often reported for mammals and birds, where genome repeat content and composition tend to be relatively stable across closely related species. The broader range observed in squamates suggests that transposable element dynamics in reptiles may follow evolutionary patterns that differ from those inferred for other vertebrate lineages [40,45]. Notably, our assembly also recovered a large proportion of unclassified repeats (25%), a feature common to other reptile genomes and likely reflecting the rapid evolution of lineage-specific transposable element families and the limited representation of reptiles in repeat libraries [40,46].

Taken together, the functional annotation and repetitive elements landscape underscore both the reliability of the predicted gene set and the strong genomic imprint of repetitive element activity on squamate protein repertoires [40,46]. The *H. orcesi* annotation provides a robust resource for future studies of adaptation and conservation and underscores the need to expand functional databases with reptile-specific data to improve annotation accuracy in this clade.

The reference genome of *H. orcesi* represents a significant step forward for conservation genomics and evolutionary research on Neotropical reptiles [47]. From a conservation perspective, this genomic resource opens the door to evaluating genetic diversity and population structure in one of Ecuador’s most threatened reptiles. These data are essential for defining evolutionarily significant units, developing science-based management plans, and monitoring genetic health over time [9]. Integrating genomic evidence into conservation actions will enable the design of strategies to preserve the adaptive potential and long-term viability of this Critically Endangered species [3].

From an evolutionary standpoint, the *H. orcesi* genome provides key insights into the molecular basis of adaptation to high-elevation, semi-arid ecosystems in the Andes. The species persists under marked seasonality, limited water availability, and high diurnal thermal variation, conditions that impose significant physiological and genetic challenges [48]. Genomic analyses will help test hypotheses on how selection has influenced metabolism, thermoregulation, and cellular stress responses in reptiles inhabiting montane dry valleys [49], a biome that remains largely unexplored at the genomic level. As one of the few high-altitude representatives of a mainly lowland lineage, *H. orcesi* offers a valuable model to investigate the genomic signatures associated with ecological transitions across altitude gradients in tropical lineages.

The genome of *H. orcesi* opens new opportunities for integrative research that combines evolution, ecology, and conservation. A key next priority will be to identify candidate genes and regulatory pathways associated with high-altitude adaptation, particularly those linked to oxygen transport, energy metabolism, and UV damage repair. Comparative analyses with lowland congeners and other teiids will help distinguish lineage-specific adaptations from broader reptilian genomic patterns, providing insight into how environmental pressures have driven diversification within the group [40]. Population genomic and landscape genetic approaches are also needed to evaluate connectivity and demographic trends among remnant populations. These analyses will inform conservation priorities by identifying barriers to gene flow, estimating effective population sizes, and assessing the genetic consequences of habitat loss and other anthropogenic pressures [50]. Integrating genomic, ecological, and spatial data will strengthen our understanding of Andean evolutionary dynamics and guide the development of evidence-based strategies for conserving *H. orcesi* and other reptiles inhabiting fragile montane dry ecosystems.

## 5. Conclusions

The genome assembly of *Holcosus orcesi* represents a major advance for the study and conservation of Ecuador’s endemic reptiles. Its high contiguity and completeness establish it as a reference-quality resource for future genomic investigations. Structural and functional annotations are consistent with those reported for other squamate genomes, further supporting the reliability of the assembly and the accuracy of the annotations.

This genomic resource opens opportunities to assess genetic diversity, population connectivity, and adaptive capacity in *H. orcesi*, a species with a critically restricted range and high susceptibility to habitat change. It also provides a platform for exploring the molecular mechanisms underlying adaptation to high-altitude and semi-arid Andean environments, which remain underrepresented in vertebrate genomics.

Integrating this reference genome with population-level sequencing will be essential for identifying candidate genes associated with thermoregulation, hypoxia tolerance, and cellular stress response.

Comparative analyses within the Teiidae family will help clarify evolutionary trajectories across altitudinal gradients, while conservation genomics applications will inform management strategies to ensure the long-term persistence of *H. orcesi*. This genome stands as both a scientific milestone and a practical tool for safeguarding Ecuador’s unique Andean biodiversity.

## Acknowledgements

We thank Wilson Ochoa for his invaluable help in locating the specimen of *Holcosus orcesi*, Renato Morales for his assistance in transporting the specimen from Cuenca to Quito, Carolina Reyes-Puig and Emilia Peñaherrera for their assistance at the Laboratory of Terrestrial Zoology and Museum of Zoology at USFQ, and Carolina Sáenz, TUERI Wildlife Hospital, for collecting the blood sample. We thank Universidad San Francisco de Quito (USFQ) for institutional support and the members of the Plant Biotechnology Laboratory (USFQ) for their support in DNA extraction, sequencing, and bioinformatics analyses.

Research activities were conducted under authorization from the Ministerio del Ambiente, Agua y Transición Ecológica del Ecuador (research permit No. MAATE-ARSFC-2025-0144 and framework contract for access to genetic resources No. MAATE-DBI-CM-2023-0313).

## Data availability

The raw sequencing reads generated for this study have been deposited in the NCBI Sequence Read Archive (SRR36082360). The genome assembly has also been uploaded to NCBI under the BioProject accession number PRJNA1153955 and BioSample accession number SAMN53165835.

